# Host bioenergetic parameters reveal cytotoxicity of anti-tuberculosis drugs undetected using conventional viability assays

**DOI:** 10.1101/2021.05.06.443046

**Authors:** Bridgette M. Cumming, Zainab Baig, Kelvin W. Addicott, D Chen, AJC Steyn

## Abstract

High attrition rates in tuberculosis (TB) drug development have been largely attributed to safety, which is likely due to the use of endpoint assays measuring cell viability to detect drug cytotoxicity. In drug development of cancer, metabolic and neurological disorders, and antibiotics, cytotoxicity is increasingly being assessed using extracellular flux (XF) analysis, which measures cellular bioenergetic metabolism in real-time. Here, we adopt the XF platform to investigate the cytotoxicity of drugs currently used in TB treatment on the bioenergetic metabolism of HepG2 cells, THP-1 macrophages, and human monocyte derived macrophages (hMDM). We found that the XF analysis reveals earlier drug-induced effects on the cells’ bioenergetic metabolism prior to cell death, measured by conventional viability assays. Furthermore, each cell type has a distinct response to drug treatment, suggesting that more than one cell type should be considered to examine cytotoxicity in TB drug development. Interestingly, chemically unrelated drugs with different modes of action on *Mycobacterium tuberculosis* have similar effects on the bioenergetic parameters of the cells, thus, discouraging the prediction of potential cytotoxicity based on chemical structure and mode of action of new chemical entities. The clustering of the drug-induced effects on the hMDM bioenergetic parameters are reflected in the clustering of the effects of the drugs on cytokine production in hMDMs, demonstrating concurrence between the effects of the drugs on the metabolism and functioning of the macrophages. These findings can be used as a benchmark to establish XF analysis as a new tool to assay cytotoxicity in TB drug development.

## Introduction

Although curative treatment is available for tuberculosis (TB), it requires adherence to a prolonged duration of drug therapy and exposes patients to drug induced toxicities. The standard treatment for drug susceptible TB includes rifampicin, isoniazid, ethambutol, and pyrazinamide over a two-month intensive phase followed by a continuation phase of rifampicin and isoniazid for 4 months. Toxicities of anti-TB drugs have been reported in up to 80% of TB patients (1), of which the most common is hepatic toxicity associated with isoniazid, rifampicin and pyrazinamide (2) in 2% to 28% of TB patients (3). Peripheral neuropathy from isoniazid, occurs in up to 40% of TB patients (4), and more frequently among TB-HIV co-infected patients (5, 6). Ethambutol has commonly been reported to cause ocular toxicity, demonstrated by either optic or retrobulbar neuritis that is either reversible or irreversible (7–9). However, medications used to treat multidrug resistant TB (MDR-TB), have much worse side effects (10). The most common adverse effects of the aminoglycosides include ototoxicity, vestibular toxicity (11, 12), nephrotoxicity and electrolyte abnormalities (13, 14). Central nervous system adverse effects have been described in adults treated with fluoroquinolones (15, 16) in particular, with cycloserine (17, 18). Furthermore, peripheral neuropathy is well documented among patients on prolonged linezolid treatment(19), including toxic optic neuropathy (20, 21). Gastrointestinal intolerance and reversible hypothyroidism are known adverse effects of ethionamide and prothionamide during long term therapy (22–24). This plethora of adverse effects is likely because most of the anti-TB drugs were discovered in the 1950’s and 1960’s with new drugs only being discovered in the last 10 years (25). The side-effects of anti-TB drugs impact adherence to chemotherapy and often result in relapse, treatment suspension or failure and development of drug-resistance. Thus, the development of new drugs with less cytotoxicity, shorter regimens and fewer side effects are needed.

Although methodologies used to assess drug cytotoxicity in the early stages of drug development have evolved over the last few years, documented “cytotoxicity” is highly dependent on the type of assay and the cell type used to assay drug toxicity. These inconsistencies urge us to ask what defines cytotoxicity, what should be measured and how can it be measured? Most standard cytotoxicity assays used in anti-TB drug development are endpoint assays measuring the viability or the integrity of the membranes of the cells. Viability assays include different classes of colorimetric tetrazolium reduction, resazurin reduction, protease markers and ATP detection (26). Endpoint assays that assess the integrity of the cell membrane include the lactate dehydrogenase release assay and trypan blue exclusion assays. However, these assays only measure how the drug affects one parameter of the cell, of which not all accurately represent the onset of cytotoxicity. Furthermore, these endpoint assays do not allude to alterations in the health of the cell that could potentially impact the functions of the cells in the absence of death. Nonetheless, defining the health status of a cell proves to be challenging in that there is no clear delineation as to what we measure and how we measure it.

High attrition rates in recent drug development have been ascribed to safety issues with organ toxicity (27, 28), which has subsequently been proven to be due to or has strong evidence suggesting links to mitochondrial impairment (29–34). This has led to the development of several *in vitro* assays to measure mitochondrial function. Measurement of the oxygen consumption rate (OCR) in real-time gives an indirect measurement of mitochondrial respiration. The Extracellular flux (XF) analyzer (Agilent) enables high-resolution, real-time multi-well plate readings of OCR in addition to real time measurements of changes in the extracellular proton concentration to provide extracellular acidification rate (ECAR), which is considered an indirect measure of glycolysis. The response of OCR and ECAR to the consecutive addition of known mitochondrial and electron transport chain (ETC) modulators or stressors is used to calculate bioenergetic parameters associated with OXPHOS and metabolism of the cells, namely, basal respiration, basal ECAR, ATP-linked OCR, compensatory ECAR, maximal respiration, spare respiratory capacity, proton leak and non-mitochondrial respiration (35). In some cases, these parameters can reveal drug cytotoxicity that is not detected in measurements of oxygen consumption rate (OCR) or extracellular acidification rate (ECAR) alone (36). ATP-linked OCR, determined from the decrease in basal respiration after the addition of an inhibitor of ATP-synthase (Complex V), oligomycin, has also been used to identify drugs that induce mitochondrial toxicity by reducing, or inhibiting the activity of ATP synthase (37). Uncoupled respiration, induced by ionophores, results in maximal respiration that has been used as an indicator of the integrity of the ETC after drug treatment (38, 39). Extracellular flux analysis, has been used to assess the cytotoxicity of drugs (30, 40–42) in the treatment of depression (43), the cancer field (44), anesthetics (45), antibiotics (39, 46) and metabolic disorders (38). However, extracellular flux analysis has not yet been investigated as a potential platform to identify cytotoxic insults induced by anti-TB drugs.

Cell lines most often used to test the cytotoxicity of anti-TB drugs and new TB drug leads include, the human hepatocellular carcinoma cell line, HepG2 cells, to assess the hepatotoxicity of the drugs (47, 48), the human alveolar epithelial cell type 2 carcinoma cell line A549 (49–51) as the lung epithelium is the first lung surface coming into contact with *Mycobacterium tuberculosis* (*Mtb*) (52), THP-1 human monocytic cell line (53, 54) and Vero (African green monkey kidney epithelial carcinoma) cells (55). Here, we adopted extracellular flux analysis as a rapid real-time platform to investigate the cytotoxicity of nine anti-TB drugs individually, and in combinations, currently used to treat drug susceptible and multi-drug resistant TB on three human cell types, the HepG2 hepatocyte cell line, phorbol myristate acetate differentiated THP-1 monocytes, and human monocyte derived macrophages (hMDM). Eight bioenergetic parameters calculated from the extracellular flux assay were compared to the viability results obtained from a MTT (3-(4,5-Dimethylthiazol-2-Yl)-2,5-Diphenyltetrazolium Bromide) tetrazolium reduction assay. This assay measures the ability of NAD(P)H dependent oxidoreductase enzymes to reduce a tetrazolium salt to formazan and is considered a measure of metabolism (56). These modulations of the bioenergetic parameters were compared with the effects of the anti-TB drugs on the functions of the hMDMs by measuring the cytokine levels in the supernatants of the hMDMs following treatment with the anti-TB drugs.

## Results

### Experimental design

We explored the potential of extracellular flux analysis as a platform to assess the cytotoxic effects of anti-TB drugs on the energy metabolism of human cells according to the workflow diagram in Fig. 1. Three cell types (HepG2, THP-1 macrophages and hMDMs) were treated with anti-TB drugs individually or in combination for 24 hours. The effects of the anti-TB drugs on the cells were analyzed by (1) extracellular flux analysis to determine eight bioenergetic parameters, (2) the MTT assay to determine viability, and (3) in the case of the drug-treated hMDMs, the culture supernatant was collected for cytokine analysis. The resultant bioenergetic parameters, viabilities and cytokine production of the drug-treated cells were analyzed using hierarchical clustering, Pearson’s correlation co-efficient and principal component analysis (PCA).

**Figure 1.**
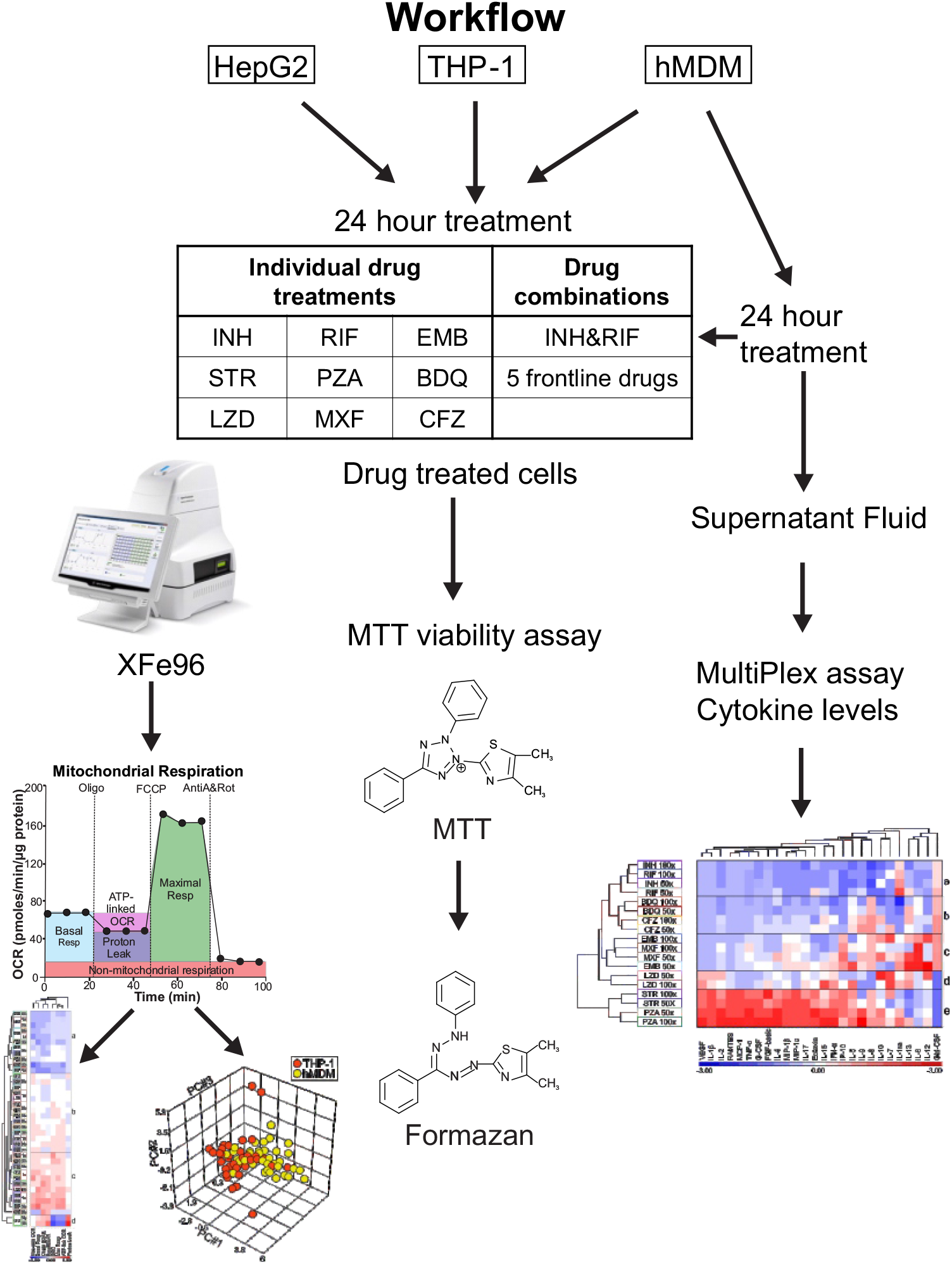
Workflow to assess extracellular flux analysis as a platform to assess the cytotoxicity of anti-TB drugs.

The three cell types, HepG2 (hepatocytes), phorbol myristate acetate differentiated THP-1 monocytes (THP-1) and human monocyte derived macrophages (hMDM) were treated with nine anti-TB drugs individually or two drug combinations for 24 hours (Fig. 1). HepG2 cells are a human hepatoma cell line that is commonly used to investigate the hepatic metabolism and hepatic toxicity of new drug leads that is induced by mitochondrial dysfunction resulting from the drugs directly targeting the electron transport chain (57). Thus, we compared the effects of the anti-TB drugs on the bioenergetics of the HepG2 cells with that of two human macrophage cell types, human monocyte derived macrophages (hMDMs) and a human THP-1 monocytic cell line that is differentiated with phorbol 12-myristate 13-acetate (PMA) to macrophages. We chose terminally differentiated human macrophages as these are usually the first immune cells to come into contact with *Mtb* through the aerosolized route of infection. We did not use mouse macrophages, due to the reported gene-expression and metabolic differences between human and murine macrophages (58–61). As larger deviations are observed in the response of hMDMs derived from different donors, responses of macrophages generated from the terminal differentiation of the human THP-1 monocytic cell line were also investigated.

The cells were treated with 1x, 10x, 50x and 100x the minimum inhibitory concentration (MIC) of the drug against *Mtb*, in the case of isoniazid (INH), rifampicin (RIF), pyrazinamide (PZA), ethambutol (EMB), moxifloxacin (MXF), clofazimine (CFZ) and linezolid (LZD); MIC_50_ in the case of BDQ and MIC_10_ in the case of streptomycin (STR). The MIC concentrations were used to enable comparisons among the effects of the different drugs on three cell types, as the physiological concentrations of the drugs vary between the serum and site of infection, in addition to differing protein binding capacities of the drugs and variable drug absorption, metabolism, distribution and perfusion of the infected areas among TB patients. As TB chemotherapy requires combination therapy to prevent the development of *Mtb* drug-resistance and to combat the tendency of *Mtb* to persist in the face of drug treatment (62, 63), two sets of drug combinations were also examined at 1x and 10x MIC of the drugs in the combination. The first combination included all the drugs used in frontline treatment of drug susceptible TB: INH, RIF, PZA, EMB and STR (64). The second combination just included INH and RIF as TB patients are treated with these two drugs for four months of their six-month regimen.

### Bioenergetic parameters derived from extracellular flux analysis

In the extracellular flux analysis, we used the Cell Mito Stress Test (CMST) on the XFe96 to calculate eight bioenergetic parameters of the untreated and drug-treated cells (Fig. 1). During the CMST run on the XFe96, mitochondrial modulators are added to the cells and the resulting changes in oxygen consumption rate (OCR) and extracellular acidification rate (ECAR) are used to calculate the following bioenergetic parameters: the basal respiration (Basal Resp), ATP-linked OCR, proton leak, maximal respiration (Max Resp), spare respiratory capacity (SRC), non-mitochondrial respiration (Non-mito OCR), basal extracellular acidification rate (Basal ECAR), and compensatory extracellular acidification rate (Comp ECAR) (Fig. 2A-C) (35).

**Figure 2.**
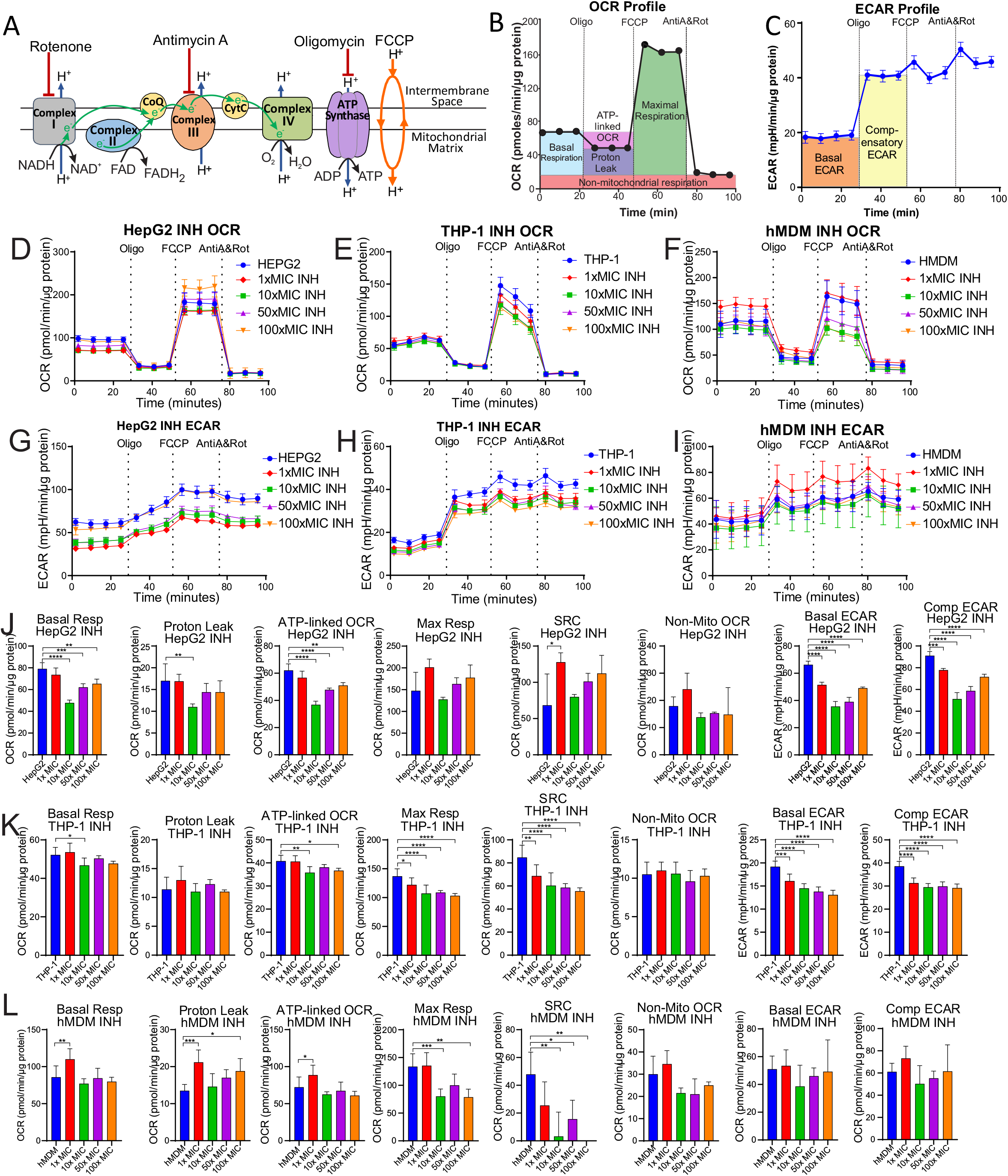
Bioenergetic parameters are calculated from the OCR and ECAR XF profiles of the CMST assay. (A) Modulators of the mitochondrial electron transport chain used to determine the ioenergetic parameters. (B) CMST profile demonstrating measurement of associated parameters of mitochondrial respiration. (C) ECAR profile demonstrating how basal ECAR and compensatory ECAR are measured from the CMST assay. (D-F) Representative CMST profiles and (G-I) ECAR profiles of HepG2 cells (D, G), THP-1 cells (E, H) and hMDMs (F, I) treated with 1x, 10x, 50x and 100x MIC INH for 24 hrs. (J-L) Bioenergetic parameters of the (J) HepG2 cells, (K) THP-1 cells, (L), hMDM cells treated with increasing MIC of INH for 24 hrs calculated from the representative profiles in (D-I).

Initially in the CMST, the respiration (OCR) of the untreated or drug-treated cells are measured to determine the basal respiration (Fig. 2B). This is followed by the addition of oligomycin, which inhibits complex V (ATP synthase, Fig. 2A), to establish how much oxygen is consumed in the production of mitochondrial ATP by complex V. This ATP-linked OCR is equivalent to the decrease in the OCR following the addition of oligomycin (Fig. 2B). Subsequently, an ionophore, carbonyl cyanide 4-(trifluoromethoxy) phenylhydrazone (FCCP) is added to the cells, which allows protons to leak across the mitochondrial membrane into the matrix. This depolarizes the mitochondrial membrane potential resulting in the ramping up of electron transport in the electron transport chain (ETC) to pump protons out of the matrix to re-establish the proton gradient and the mitochondrial membrane potential. The increased electron transport increases the oxygen consumption at complex IV and enables a measurement of the maximal respiration of the cell (Fig. 2A and B). After the addition of antimycin A and rotenone, inhibitors of complex III and complex I, the ETC is shut down resulting in inhibition of mitochondrial OCR, and the resultant OCR gives a measurement of the non-mitochondrial respiration (Fig. 2A and B). This non-mitochondrial OCR is then subtracted from the basal respiration and the maximal respiration to give their true mitochondrial values. The difference between the non-mitochondrial respiration and the ATP-linked OCR is equivalent to the proton leak, which is the measure of oxygen consumption at Complex IV that is not linked to ATP production and involved in restoring the mitochondrial membrane potential that is depolarized by the natural leak of protons into the mitochondrial matrix. SRC, which gives a measurement of the cell’s capacity to respire under conditions of stress, is calculated by the difference between the maximal respiration and the basal respiration. Extracellular acidification is generated by lactic acid that is derived from pyruvate, the end-product of glycolysis, and carbonic acid that is derived from carbon dioxide produced by the TCA cycle (65). ECAR measurements made simultaneously during the CMST run are used to determine the basal ECAR (ECAR prior to the addition of oligomycin) and compensatory ECAR (after the addition of oligomycin). Compensatory ECAR is induced by the inhibition of mitochondrial ATP synthase with oligomycin, which results in increased glycolysis to generate ATP for the cell’s demands (Fig. 2B).

Representative CMST OCR and ECAR profiles of each cell type are illustrated in Fig. 2D-I together with the altered profiles of the cells after treatment with increasing concentrations of isoniazid (INH). The calculated bioenergetic parameters from the HepG2 profiles in Fig 2D and 2G are shown in the Fig. 2J panel; from the THP-1 profiles in Fig. 2E and H are shown in Fig. 2K; and from the hMDM profiles in Fig. 2F and I are shown in Fig. 2L. These bioenergetic parameters provide quantitative measurements of aspects of mitochondrial respiration, non-mitochondrial respiration, and extracellular acidification, which enable assessment of how a potential drug affects the bioenergetic pathways of the cell that provide ATP. For instance, increasing concentrations of INH decreased both the Basal Resp and the ATP-linked OCR of the HepG2 cells (Fig. 2J) and to a lesser extent in THP-1 cells (Fig. 2K), which are both essential for promoting OXPHOS. In both macrophage models, THP-1 cells and hMDMs, INH decreased both the Max Resp and the SRC that both give a measure of the cell’s ability to respond to conditions of stress (Fig. 2K and 2L). In hMDMs, INH increased the proton leak, which results in incomplete coupling of oxygen consumption and ATP synthesis in OXPHOS. Furthermore, INH decreased the basal ECAR and compensatory ECAR of both the HepG2 cells and the THP-1 cells (Fig. 2J and K) suggesting additional suppression of glycolysis. These findings suggest that INH depresses the bioenergetic pathways, particularly in the HepG2 cells and THP-1 cells.

However, moxifloxacin, for example, altered the bioenergetic parameters of these cell types differently (Fig. S1). Although increasing concentrations of moxifloxacin (MXF) reduced the Basal Resp, Max Resp and ATP-linked OCR of the HepG2 cells and the hMDMs, it had no effect on these parameters in the THP-1 cells (Fig. S1B-D). Furthermore, MXF significantly reduced the Basal ECAR and Comp ECAR in all three cell types. The bioenergetic parameters of all three cell types in response to the anti-TB drug treatments are listed in Dataset S1.

In summary, we have adopted the extracellular flux analysis platform to assess the cytotoxic effects of anti-TB drugs on the bioenergetic metabolism of HepG2, THP-1 and hMDM cells. We have used the CMST assay, which generates eight bioenergetic parameters that reveal how anti-TB drugs modulate different aspects of respiration and glycolysis. Here, we have demonstrated how increasing concentrations of two of the drugs investigated, INH and MXF altered the OCR and ECAR profiles of the three cell types and the resulting bioenergetic parameters. The responses of the bioenergetic parameters of each cell type to the drug treatment were analyzed using hierarchical clustering, Pearson’s correlation co-efficient and principal component analysis (PCA) to identify any distinct trends.

### Each cell type has a unique bioenergetic fingerprint in response to the anti-TB drugs

Hierarchical clustering was used to determine to determine if the bioenergetic metabolism of the three cell types respond similarly to the anti-TB drugs. The three cell types were treated with four concentrations of nine anti-TB drugs individually and two concentrations of two drug combinations. After 24 hrs treatment, their bioenergetic metabolism was examined using extracellular flux analysis and the CMST assay to calculate the eight bioenergetic parameters. Hierarchical cluster analysis of the bioenergetic parameters of the drug-treated cells relative to untreated cells was performed to assess if the drugs induced similar changes in the bioenergetic parameters of the different cell types. Both rows and columns of normalized relative vaues were used in the clustering where Euclidean methods were applied for disimilarities across both rows and columns. Z-normalization was perfomed to transform relative values of the bioenergetic values to avearage = 0 and SD = 1. Fig. 3 shows heat maps of the z-normalization values where rows (drugs and their concentrations) and columns (bioenergetic parameters) have been ordered based on their correlation hierarchical clustering, using the average linkage method. Hierarchical clustering of the effects of the drugs on the bioenergetic parameters of all three cell types (Fig. 3A) demonstrates no clustering according to cell type or drug. This suggested that the measurement of eight bioenergetic parameters enabled the detection of a variety of effects on the bioenergetic processes in different cells.

**Figure 3.**
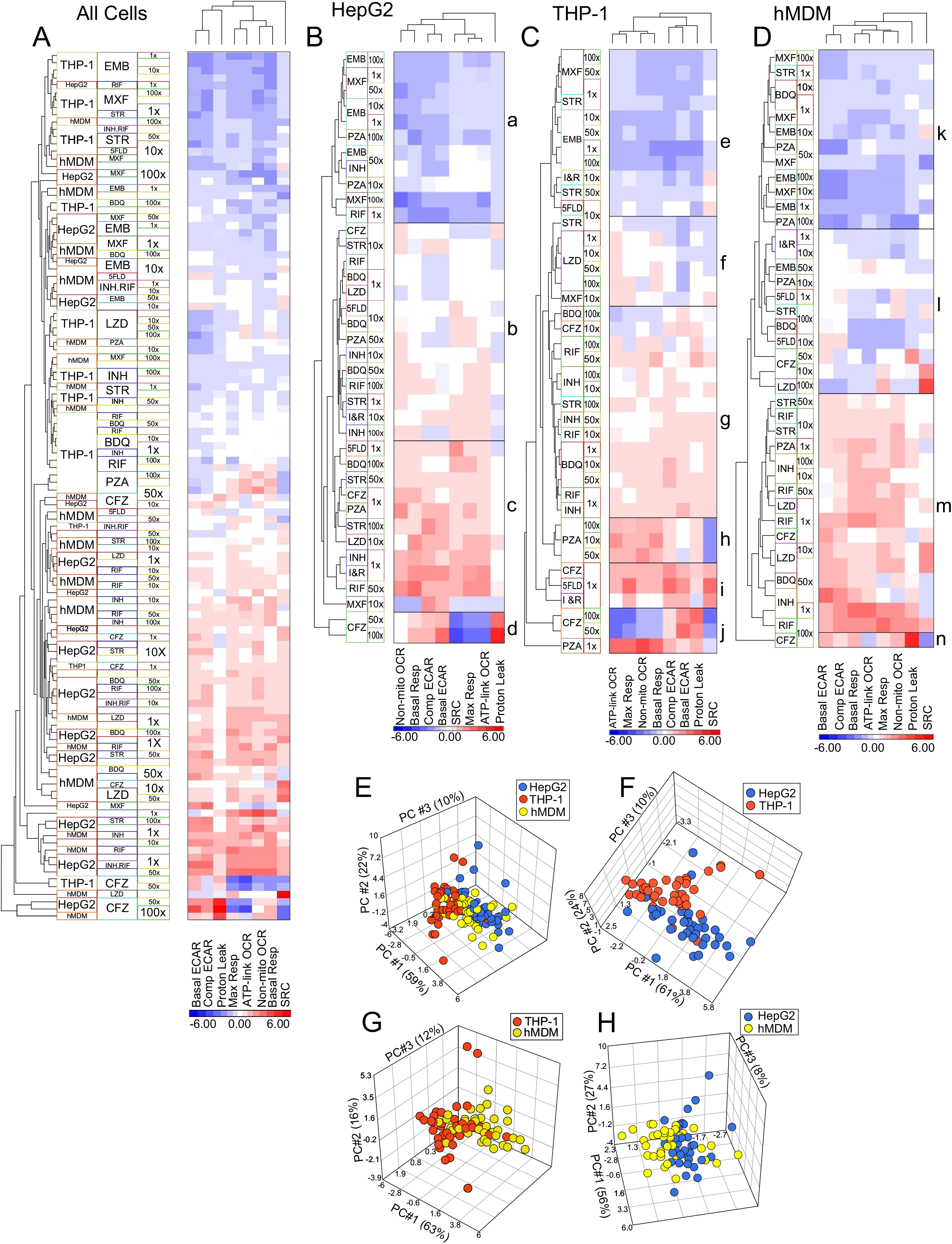
Each cell type demonstrates distinct bioenergetic fingerprints when treated with anti-TB drugs. (A-D) Heat maps of the hierarchical clustering of the z-normalization values calculated from the relative bioenergetic parameters of (A) all cell types, (B) HepG2, (C) THP-1, and (A) hMDM treated with the indicated concentrations of anti-TB drugs. (E-H) PCA analysis of the combined bioenergetic parameters of (E) all three cell types (HepG2, THP-1 and hMDM), (F) HepG2 and THP-1 cells, (G) hMDM and THP-1 cells, (H) HMDM and HepG2 cells, treated with increasing MIC concentrations of anti-TB drugs.

The heat maps of the hierarchical clustering of the effects of the drugs on the bioenergetic parameters of the individual cell types (Fig. 3B-D) demonstrate that each cell type has distinctive patterns in their response to the drug treatments. These distinct patterns of clustering observed in each cell type have been indicated with boxes to facilitate discussion. In the HepG2 cells, the effects of all four concentrations of EMB, three of MXF and two of PZA cluster together because they reduce the z-normalization values of all the bioenergetic parameters below the average (Fig. 3B, box a). The second cluster is divided into two groups with distinct patterns (Fig. 3B, box b and c). Both groups contain the same drugs, but at different concentrations. The drugs include BDQ, INH, RIF, STR, LZD, PZA, CFZ and the two drug combinations (5FLD and INH.RIF), but the concentrations of each drug are interspersed between each other. This second cluster also includes the effects of 10x MIC of MXF, but they are separated from the effects of the other drugs in the second cluster. The effects of the 50x and 100x MIC of CFZ cluster separately from the effects of all the drugs (Fig. 3B, box d) with low z-normalization values for maximal respiration, SRC and ATP-linked OCR and high z-normalization values for proton leak. In the HepG2 cells, the proton leak is distinct from the clustering of the other bioenergetic parameters, with the ATP-linked OCR and the Max Resp being the most closely linked parameters.

In the THP-1 cells, there is a great deal more clustering of the effects of several concentrations of the same drug on the bioenergetic parameters than detected in the HepG2 cells. This suggests that some of the anti-TB drugs have very distinct effects on the THP-1 cells that is not observed in the HepG2 cells. For example, the effects of all four concentrations of EMB and of LZD cluster distinctly together in Fig. 3C, box e and f, respectively. The effects induced by three concentrations each of MXF, of BDQ and of PZA cluster together distinctly (Fig. 3C, box e, f and g, respectively); and two concentrations each of INH, RIF and CFZ cluster together. However, THP-1 cells do behave similarly to the HepG2 cells in that the effects of EMB and MXF cluster together because they reduce the z-normalization values below normal (Figure 3C, box e). Yet in the case of the THP-1 cells, the effects of MXF and EMB cluster together with two STR concentrations and the 10x MIC of both drug combinations, 5FLD and INH.RIF (Fig. 3C, box e). The “block” showing the effects of the four concentrations of LZD clusters with the effects of the 10x MIC concentration of both STR and MXF (Fig 3C, box f). This cluster (box f) is closely linked to the effects of BDQ, RIF and INH clustered together with 100x MIC STR and 10x MIC CFZ, (Fig. 3C, cluster g) by inducing small or no fluctuations on z-normalization values from the average. Strikingly, the effects of three PZA concentrations are clustered on their own inducing higher than average z-normalization values in the ATP-linked OCR, Max Resp, Non-Mito OCR and Basal Resp (Fig. 3C, box h). The 1x MIC concentration of PZA has the most divergent effects on the bioenergetic parameters such that it clusters separately from all the other drugs (Fig. 3C, last row in box j). The effects of the lowest concentrations of CFZ, and both drug combinations, 5FLD and INH.RIF, cluster together (Fig 3C, box i), with the effects of the 50x and 100x MIC CFZ also clustering apart from the effects of the other drugs (Fig. 3C, box j), by inducing lower than average z-normalization values for the ATP-linked OCR, Max Resp, Non-Mito OCR and Basal Resp. The hierarchical clustering of the bioenergetic parameters indicates that the effects of the drugs on the SRC of the THP-1 cells cluster separately from the other bioenergetic parameters.

Treatment of the hMDM cells with the anti-TB drugs appears to induce more lower z-normalization values than those observed in the other two cell types (Fig. 3D). As in the case of the THP-1 cells, the effects of MXF and EMB are clustered together with the effects of 1x MIC of STR, but in the hMDMs, they are also clustered with the low concentrations of BDQ and higher concentrations of PZA because they all reduce the z-normalization values below average (Fig. 3D, box k). This cluster (box k) is closely linked to a cluster of the effects of the two drug combinations, 5FLD and INH.RIF and other concentrations of BDQ, PZA and STR that have some minimal fluctuations in the z-normalization values from average (Fig 3D, box l). The effects of 50x and 10x MIC CFZ and 100x MIC LZD cluster separately but are still linked to previously described cluster (Fig. 3D, box l). In contrast, the effects of RIF, INH, LZD, and remaining concentrations of STR, PZA and BDQ cluster together by increasing the z-normalization values of the bioenergetic parameters above average (Fig. 3D, box m). Lastly, similarly to both the HepG2 and THP-1 cells, the effects of the 100X MIC CFZ is clustered separate from the effects of all the other drugs (Fig. 3D, box n). Similar to the THP-1 cells, the drug induced changes in the SRC of the hMDMs clustered separately from the other bioenergetic parameters.

The relatedness between the cell types was examined using PCA analysis (66) of the effects of the four concentrations of the anti-TB drugs on the combined relative bioenergetic parameters of each cell type (Fig. 3E). When the PCA analysis of the cell types were compared pairwise, there was a noticeable separation between the effects of the drugs on the bioenergetic parameters of the HepG2 and THP-1 cells (Fig. 3F) that was not as well-defined when the hMDM cells were compared with the THP-1 cells (Fig. 3G) or with the HepG2 cells (Fig. 3H). This illustrates that the anti-TB drugs do not have equal effects on all cell types and other cell types should be considered when assessing cytotoxicity/modulatory effects of anti-TB drugs.

In sum, the hierarchical clustering and PCA analysis of the bioenergetic parameters generated from the extracellular flux analysis revealed that the three cell types investigated demonstrate unique bioenergetic responses to treatment with the anti-TB drugs. Hierarchical clustering demonstrated different pattens of clustering of the bioenergetic parameters of the three cell types when treated with the anti-TB drugs. In particular, PCA analysis revealed that the bioenergetic parameters of the drug-treated THP-1 cells had more noticeable separation from the HepG2 cells than the hMDM cells.

### Basal respiration correlates with all other bioenergetic parameters in all three cell types

To identify a bioenergetic parameter that correlates with the changes in the other bioenergetic parameters in response to treatment with any of the anti-TB drugs, Pearson’s correlation co-efficient was calculated for all pairwise combinations of the combined bioenergetic parameters of all three cell types combined (Fig. 4A) and for each cell type individually (Fig. 4B-D). The left panel in Fig. 4A shows a heat map of the correlation co-efficient of each pairwise comparison of the combined bioenergetic parameters for all the cells, and in the right panel, the averages of these correlation co-efficient for each bioenergetic parameter is given in a heat map to demonstrate the parameters with the highest correlations. The Pearson’s correlation co-efficient for the bioenergetic parameters of each cell type were demonstrated in a similar manner (Fig. 4B-D).

**Figure 4.**
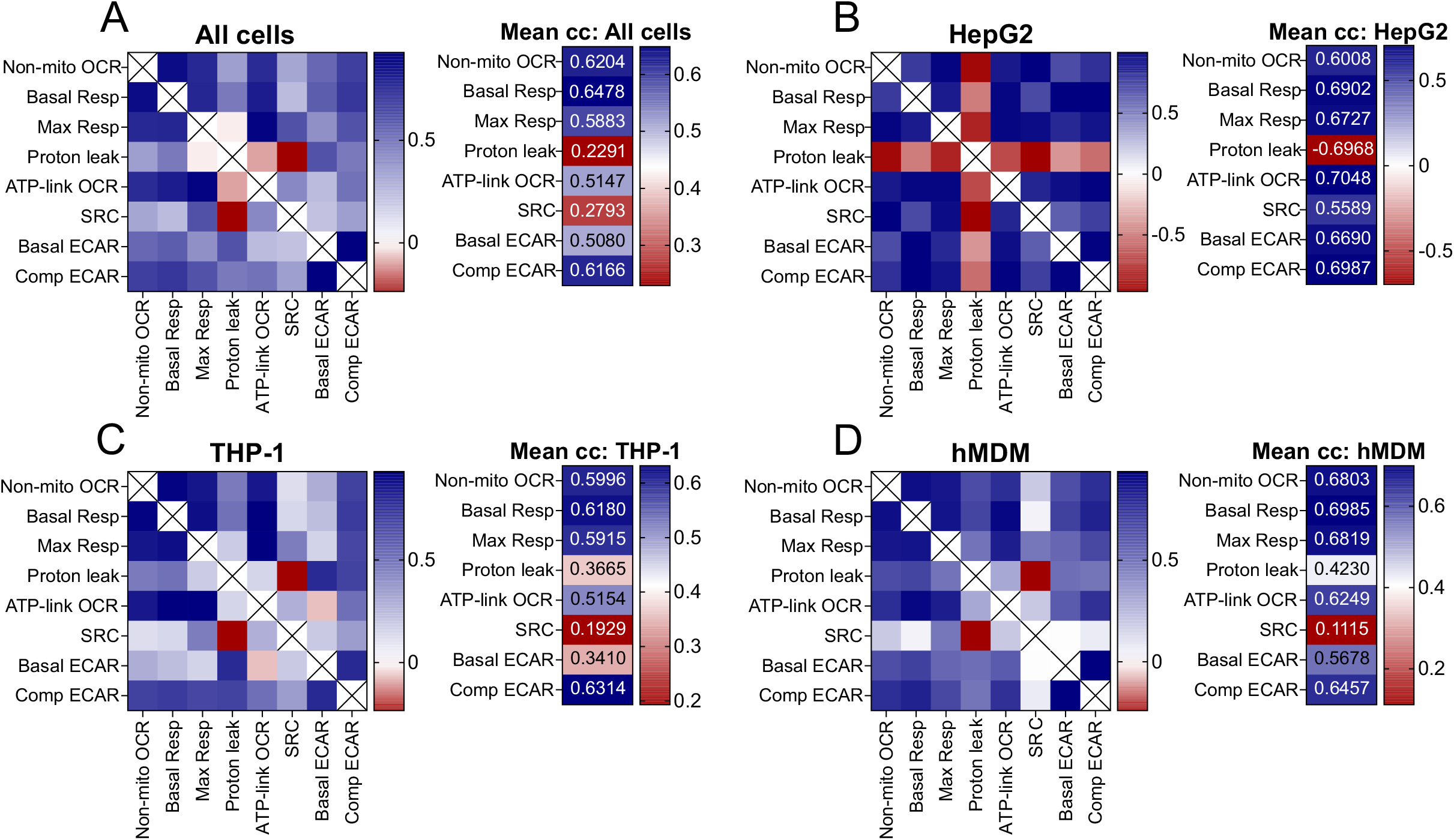
Basal respiration correlates with all the bioenergetic parameters. Heat maps of the Pearson correlation co-efficient of the averaged bioenergetic parameters induced by anti-TB drug treatment (Left panel), and the means of the correlation co-efficient (Right panel) of (A) all cell types, (B) HepG2 cells, (C) THP-1 cells and (D) hMDMs treated with increasing MIC concentrations of anti-TB drugs. (Refer to Dataset S2-S5 for the Pearson Correlation co-efficient used to plot the heat maps).

When the bioenergetic parameters of all cell types were analyzed, the relative Basal Resp had the highest correlation co-efficient (r = 0.64, Fig. 4A) with all the other bioenergetic parameters of all three cell types. In the case of the HepG2 cells, several bioenergetic parameters behave similarly to each other in response to the anti-TB drugs, with relative ATP-linked OCR and relative Comp ECAR having the highest correlation co-efficient (r = 0.70, Fig. 4B), followed by relative Basal Resp (r = 0.69). Notably, the proton leak in the HepG2 cells exhibited a strong negative correlation with all the other bioenergetic parameters. This demonstrates strong inverse relationships between proton leak and the other bioenergetic parameters of the HepG2 cells when treated by the anti-TB drugs. In THP-1 cells, the relative Comp ECAR (r = 0.63, Fig. 4C) and Basal Resp (r = 0.61) had the highest correlations with the other bioenergetic parameters in response to the anti-TB drugs. In hMDM cells, relative Basal Resp correlated with the changes in the other respiratory parameters in response to the anti-TB drugs (r = 0.70, Fig. 4D) with Max Resp and Non-mito OCR also having a high correlation (0.68 in both cases).

Conversely, the bioenergetic parameter with the lowest correlation with the other bioenergetic parameters was proton leak in HepG2 cells (r = −0.70, Fig. 4B), SRC in THP-1 cells (r = 0.20, Fig. 4C), and SRC in hMDMs (r = 0.11, Fig. 4D). When the bioenergetic parameters of all the cells were combined, proton leak had the lowest correlation (r = 0.23, Fig. 4A), followed by the SRC (r = 0.30). This suggests that SRC and proton leak respond differently from the other parameters to anti-TB drug treatment, possibly due to increased sensitivity. This is feasible with SRC, given that SRC is a measure of the ability of the cell to respire under conditions of stress, which may be compromised with the anti-TB drug treatment.

Overall, we conclude that Basal Resp can be used to assess the overall changes in the bioenergetic metabolism induced by the anti-TB drugs as it has a high correlation co-efficient with the effects of the anti-TB drugs on the other bioenergetic parameters amongst all three cell types. Furthermore, Basal Resp has the highest correlation co-efficient in hMDMs. Another parameter that demonstrated high correlation with the other bioenergetic parameters in all three cell types was Comp ECAR, with it being the parameter with the highest correlation in THP-1 cells. In HepG2 cells, which are more oxidative than glycolytic, ATP-linked OCR is the parameter that has the highest correlation with the other bioenergetic parameters. SRC and proton leak demonstrated the least correlation with the other bioenergetic parameters, probably in response to different mechanisms of toxicity induced by the anti-TB drugs.

### The bioenergetic parameters enable separation of distinct effects of anti-TB drugs

To identify if groups of anti-TB drugs have similar effects on the bioenergetic parameters of each cell type, the identity of the anti-TB drugs were selected in the PCA analysis on each cell type. Clustering of the drug effects on the bioenergetic parameters of the cell was most distinct in the THP-1 cells. Fig. 5A shows how the modulation of the THP-1 bioenergetic parameters induced by EMB, MXF and LZD cluster separately from the effects of RIF, INH and BDQ on the THP-1 cells. The effects induced by PZA on the bioenergetics of the THP-1 cells are separated from the EMB-MXF-LZD and RIF-INH-BDQ clusters. The effects of the highest concentrations of CFZ (50x and 100x MIC) on THP-1 bioenergetics are even further removed from the effects of the other drugs. This reflects the hierarchical clustering of the effects of the drugs on the THP-1 bioenergetic parameters observed in the heatmap in Fig. 3C.

**Figure 5.**
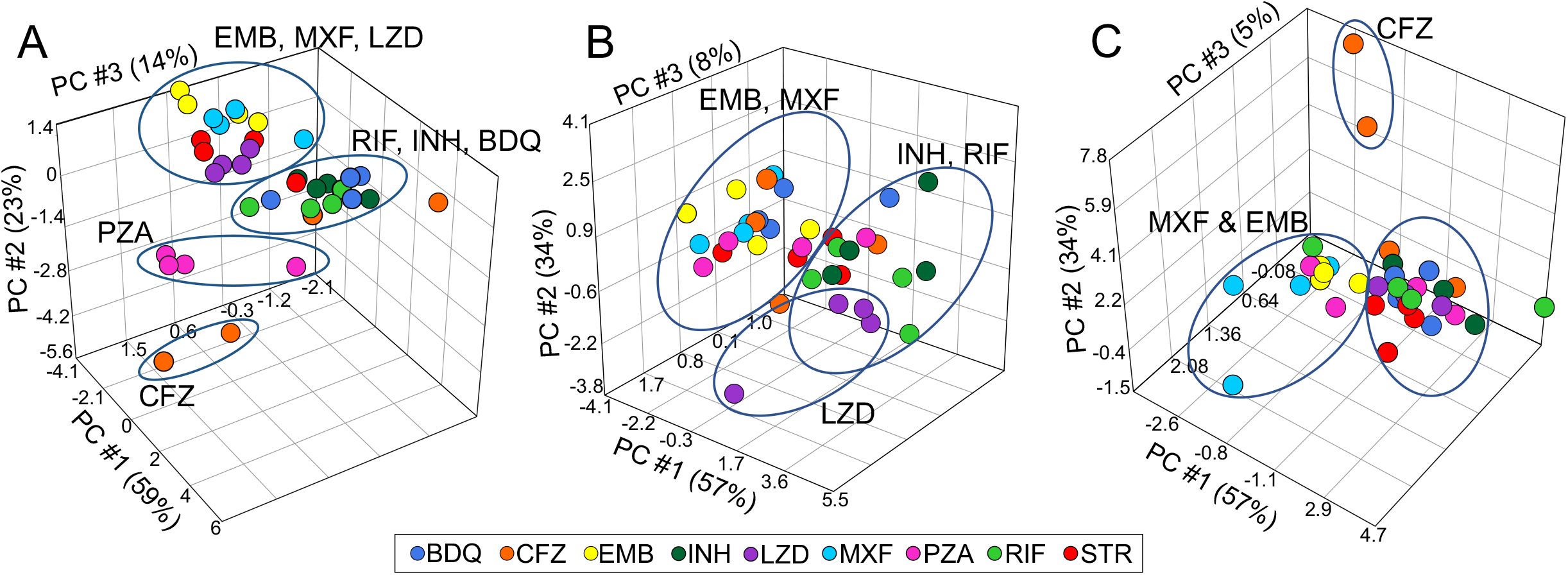
The bioenergetic parameters can be used to distinguish the effects of anti-TB drugs on the cells. (A-C) PCA analysis of the averaged bioenergetic parameters of the (A) THP-1 cells, (B) HMDM cells, and (C) HepG2 cells, that were treated with increasing concentrations of anti-TB drugs demonstrating clustering of the different drug treatments. The color of the spheres indicates different drug treatments.

In hMDM cells, the effects of EMB and MXF on the bioenergetics are again separated from the effects of INH and RIF on the hMDM bioenergetics (Fig. 5B). Specific to the hMDMs, the effects of LZD on the bioenergetics separated from the bulk of the effects of the other TB drugs. The effects of 100x MIC of CFZ was detached from the bioenergetic modulations generated by the other anti-TB drugs. This is supported by the hierarchical clustering of the effects of the drugs on the hMDM bioenergetic parameters in Fig. 3D.

In the HepG2 cells, the separation of the effects on the anti-TB drugs on the bioenergetic parameters were less defined (Fig. 5C). As in the THP-1 and hMDM cells, the bioenergetic parameters of the MXF and EMB treated HepG2 cells were separated from the bioenergetic parameters of most of the other drug-treated cells (Fig. 5C). Again, the bioenergetic modulations induced by the highest concentrations of CFZ were clearly disconnected from the effects of the other anti-TB drugs on the bioenergetics of the HepG2 cells. The clustering of MXF and EMB and the disconnection of CFZ align with the hierarchical clustering of the effects of the anti-TB drugs on the HepG2. Overall, these findings indicate that the bioenergetic parameters can be used to identify groups of drugs with similar effects on the bioenergetic metabolism of the cells.

### The bioenergetic parameters reveal a broader range of effects than % viability

As viability assays have been conventionally used to assess the cytotoxicity of new drug leads, we compared the percentage (%) viabilities obtained in MTT assays for each drug-treated cell type with the bioenergetic parameters obtained under the same conditions. We calculated the Pearson’s correlation co-efficient for all possible pairwise combinations of the bioenergetic parameters with the % viabilities. Heatmaps in Fig. 6A-D reveal that viabilities of all the drug treatments negatively correlated with all the averaged bioenergetic parameters when the parameters of all three cell types were combined and separated by cell type. These negative correlations were all below −0.4.

**Figure 6.**
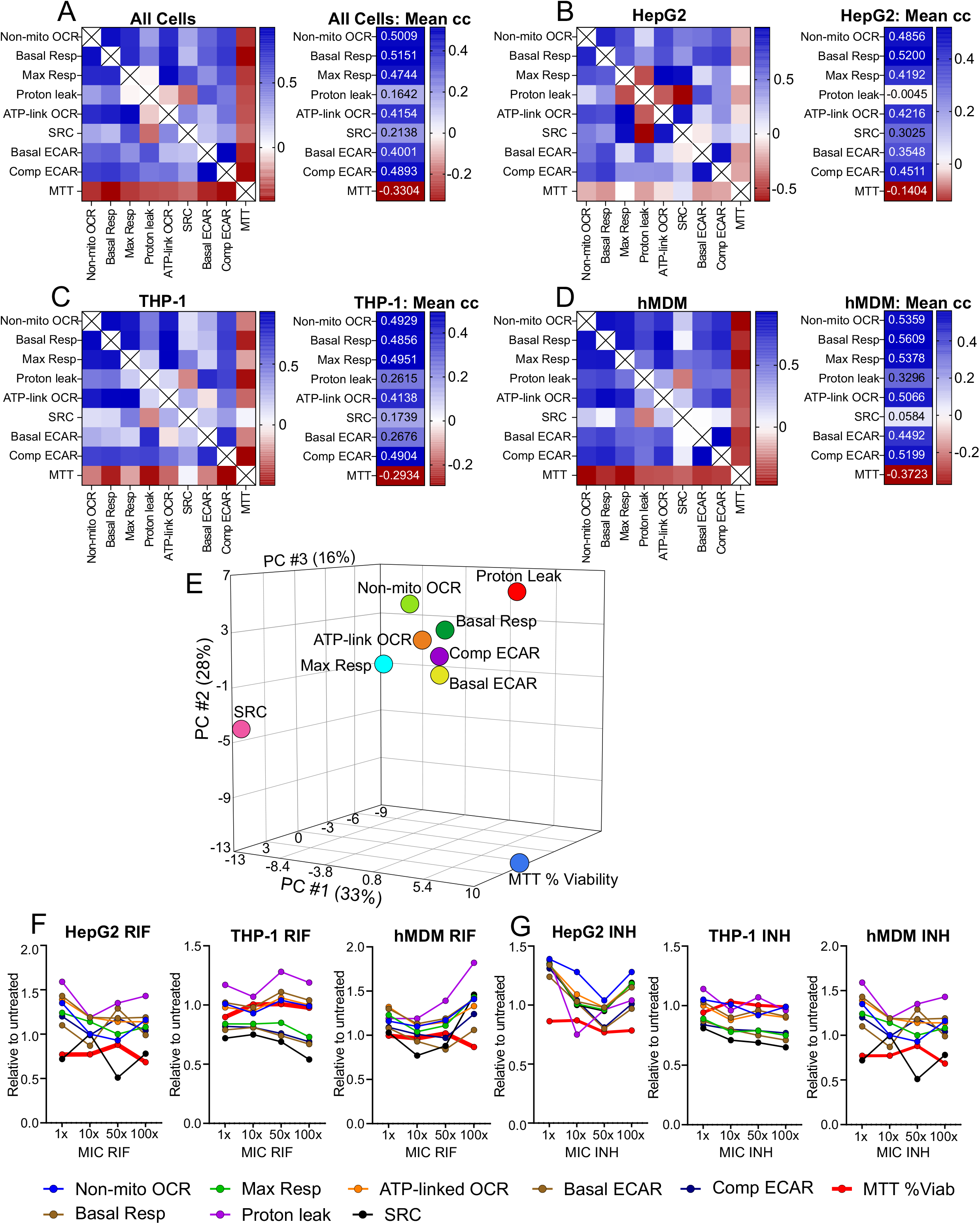
The bioenergetic parameters display a wider range of effects of the anti-TB drugs than the MTT assay. (A-D) Heat maps of the Pearson’s correlation co-efficient of MTT % viability correlated with the averaged bioenergetic parameters pairwise (left panel), and an adjacent heatmap of the means of the correlation co-efficient (right panel) of (A) all cell types, (B) hMDMs, (C) THP-1 cells, and (D) HepG2 cells treated with increasing MIC concentrations of anti-TB drugs. (Refer to Datasets S6-S9 for the values of the Pearson correlation co-efficient used to plot the heat maps). (E) PCA analysis depicts separation of the cumulative MTT %viability values from the averaged individual bioenergetic parameters. (F-K) Line plots of relative bioenergetic parameters and the MTT %Viability of the three cell types treated with increasing MIC concentrations of (F) RIF and (G) INH.

This poor correlation between the % viability and the bioenergetic parameters of the cells was supported by PCA analysis of the bioenergetic parameters and % viabilities of all the cell types combined. Fig. 6E demonstrates the clear separation of the % viabilities of the MTT assay from all the bioenergetic parameters. Furthermore, PCA analysis shows that the relative SRC is separated to the greatest degree from the other bioenergetic parameters, which are clustered to some extent. The SRC is separated to the largest degree from proton leak. This is also supported by the Pearson correlation analysis where the relative SRC has the lowest positive correlation co-efficient with the other bioenergetic parameters in both the THP-1 and hMDM cells (Fig. 6C and D). However, in HepG2 cells and in the combination of the bioenergetic parameters of all three cell types, the relative proton leak has the lowest positive correlation co-efficient, followed by SRC (Fig. 6A and B).

Line plots of the individual relative bioenergetic parameters and % viabilities versus concentrations of the drugs demonstrate variability in the parameters and % viabilities with an increase in drug concentration (Fig. S2). Fig. 5F and G shows representative line plots of the bioenergetic parameters and the % viability (thick red line) of the three cell types when treated with RIF (Fig. 5F) or with INH (Fig. 5G). The line plots demonstrate that the bioenergetic parameters of the cells exhibit a much broader range of effects than that observed in the % viability with an increase in drug concentration.

In summary, the bioenergetic parameters reveal changes in the energy metabolism of the three cell types induced by the anti-TB drugs that are not reflected by changes in the % viability measured using the MTT assay. This is due to the extracellular flux analyzer being an extremely sensitive instrument in that it measures oxygen consumption in pmoles O_2_/min and extracellular acidification in mpH/minute in addition to generating eight bioenergetic parameters with the addition of metabolic modulators. In comparison, the MTT assay measures the activity of one metabolic enzyme, nicotinamide adenine dinucleotide phosphate (NADPH)-dependent cellular oxidoreductase, in the cells as a measure of the metabolic activity and viability. This suggests that the bioenergetic parameters of the cell measured using extracellular flux analysis are more sensitive than the MTT assay.

### Clustering of the effects of anti-TB drugs on hMDM cytokine production resembles that observed in the hMDM bioenergetic parameters

As metabolism has been demonstrated to be intricately related to immunity(67), we investigated the effects of the anti-TB drugs on the cytokine production of the hMDMs. To determine the effects of the anti-TB drugs on the function of the macrophages, the cytokine production of hMDMs after treatment with the two highest concentrations of each drug was assessed in a multiplex assay. Hierarchical cluster analysis of the cytokine production of the drug-treated hMDMs relative to the untreated hMDMs was performed. Fig. 7 shows a heat map of the z-normalization values where rows (drugs and their concentrations) and columns (cytokines) have been clustered based on their correlation hierarchical clustering or similarities, using the average linkage method.

**Figure 7.**
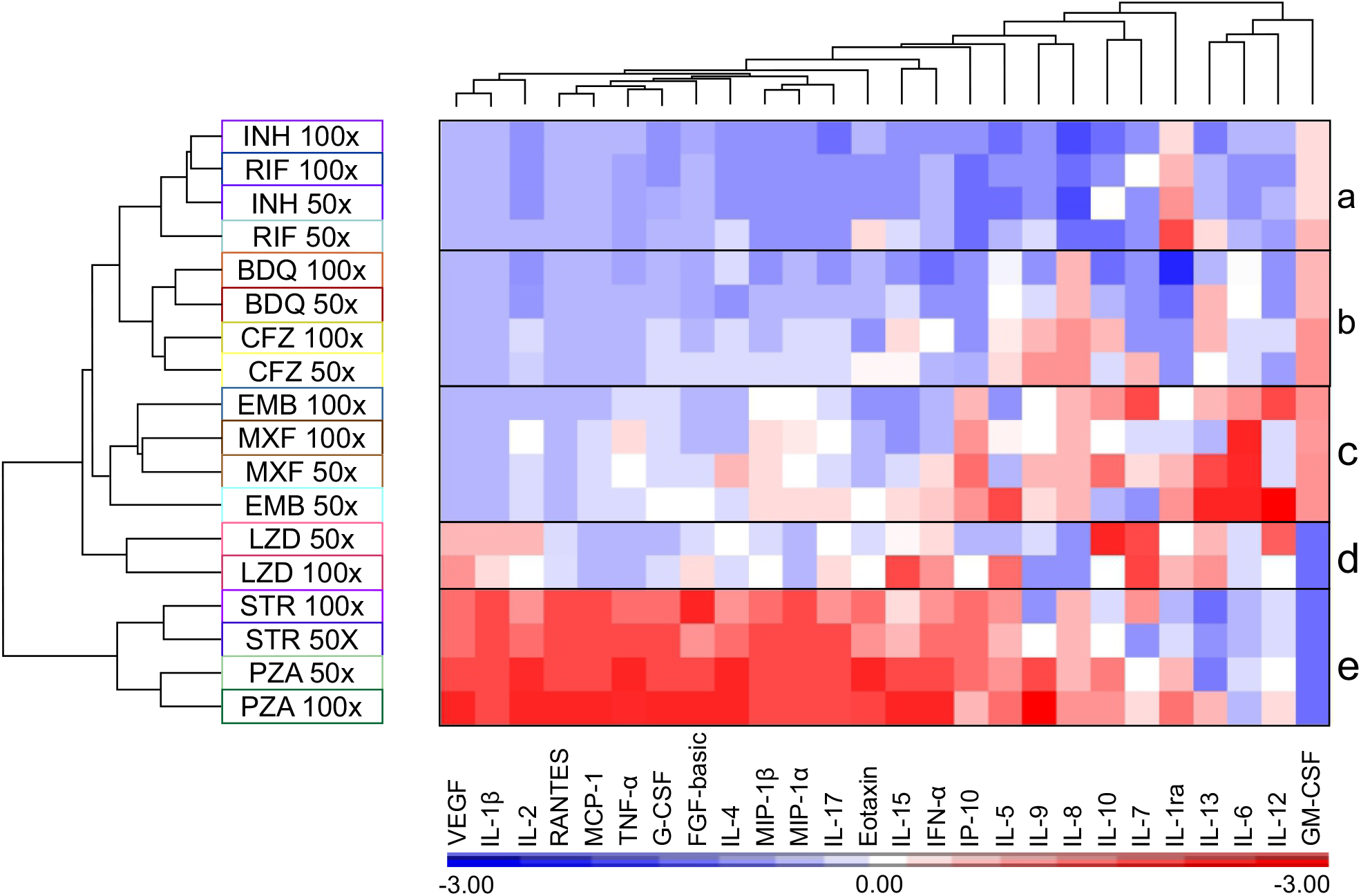
The anti-TB drugs alter cytokine production of the hMDM cells in a pattern resembling that of the hMDM bioenergetic parameters. Heatmap and hierarchical clustering of the z-normalization values of the cytokines produced by hMDMs after treatment with the indicated concentrations of the anti-TB drugs.

As observed in the hierarchical clustering of the bioenergetic parameters (Fig. 3D) and the PCA analysis (Fig. 5B) of the hMDMs treated with anti-TB drugs, the effects of the INH and RIF on the cytokine production clustered together reducing the z-normalization values of the cytokine production below the average of all the drugs investigated (Fig. 7, box a). The only exceptions are the z-normalization values of IL-1ra and GM-CSF, which were increased above average in the case of IL-1ra with only slight increments in GM-CSF. IL-1ra is the IL-1 receptor antagonist and it inhibits the pro-inflammatory action of IL-1 (68). Linked to the effects of INH and RIF, are the effects of BDQ and CFZ that cluster together (Fig. 7, box b). Likewise, BDQ and CFZ also reduce the z-normalization values of most of the cytokines below the average. Cytokines that were increased above the average include, IL-8 and GM-CSF by both CFZ and BDQ, IL-9 by CFZ, and slight increases in IL-13. GM-CSF stimulates the differentiation and proliferation of myeloid progenitors in the bone marrow, and induces the activation and migration of myeloid progenitors to sites of inflammation (69). IL-8 is a chemokine that attracts neutrophils to regions of inflammation and activates them. Both IL-9 and IL-13 are Th2 cytokines that inhibit the pro-inflammatory response (70, 71), but IL-9 promotes mast cell growth and function (72) and IL-13 mediates allergic inflammation and asthma^5^.

These two clusters mentioned above are linked to the clustered effects of EMB and MXF on the hMDM cytokine production. These drugs induce both a decrease and an increase the z-normalization values above average, with a particularly high induction of the z-normalization values of IL-6 and IL-13. EMB also increases the values of IL-12, a pro-inflammatory cytokine (Fig. 7, box c). The clustering of cytokine production resulting from treatment of the hMDM cells with EMB and MXF is supported by the hierarchical clustering of the effects of EMB and MXF on the bioenergetic parameters of the hMDM cells (Fig. 3D, box d) and the clustering observed in the PCA analysis of the effects of MXF and EMB on the bioenergetic parameters of the hMDM (Fig. 5B). LZD clusters on its own because it increases the z-normalization values of different cytokines, in particular IL-7 and IL-10 at 50x MIC, with slight increases in VEGF and IL-1β (Fig. 7, box d). This also correlates with the distinct clustering of the effects of all four concentrations of LZD on the bioenergetic parameters of the hMDM cells in the PCA analysis in Fig. 5B apart from the effects of the other drugs. In contrast to the previous drugs, it reduces the z-normalization values of GM-CSF. Lastly, STR and PZA cluster together with a striking inverse of the patterns of the z-normalization values observed in the previous clusters (Fig. 7, box e). These cytokines with high z-normalization values include pro-inflammatory cytokines such as IL-1β, IL-2, TNF-α, IFN-γ, IL-17 and chemokines such as MCP-1, MIP-1α, MIP-1β and IP-10. This suggests that STR and PZA potentially induce pro-inflammatory modulation of the macrophages. However, like LZD, STR and PZA also reduce the z-normalization values of GM-CSF.

In summary, the hierarchical clustering of the cytokines produced by the anti-TB drug treated hMDM cells resemble the clustering observed in the bioenergetic parameters of the drug treated hMDMs in the PCA analysis, especially in the case of RIF, INH, EMB, MXF and LZD. Interestingly, treatment of the hMDM cells with STR or PZA induce a completely different pattern of cytokine production to the other anti-TB drugs, suggesting induction of a pro-inflammatory response.

## Discussion

High attrition rates in drug development during the clinical and post-market phases due to safety issues underscore the need for new technology to screen for “cytotoxicity” at early stages in drug development. In anti-TB drug development, current methods used to assess cytotoxicity of new chemical entities have a single endpoint measurement as an indicator of cell viability, which does not reflect earlier events of distress induced by the compounds in the absence of cell death. Here, we adopted a multi-well non-invasive extracellular flux analysis platform that rapidly detects the modulation of bioenergetic metabolism of cells induced by anti-TB drugs in real-time prior to cell death that is measured by conventional viability assays. This rapid detection of earlier events of anti-TB drug induced bioenergetic distress provides a novel tool to detect early toxicity affecting the health of the cells that potentially leads to cellular dysfunction, thereby reducing high attrition and costs involved in further drug development.

The MTT assay, and other tretrazolium reduction assays have often been used to assess the cytotoxicity of anti-TB drugs, new chemical entities, or combinations (48, 50, 73–75). However, the major drawback of viability assays is that they only focus on the effects of the drugs on one aspect of metabolism, such as the generation of oxidized reducing equivalents, contributing to the viability of the cells and are not sensitive enough to detect alterations to the health of the cell that would impact the functioning of the cell in the absence of cell death. Yet, it is not clear what parameters define the health of the eukaryotic cell nor how they can be measured. As energy in the form of ATP is required by all eukaryotic cells to survive and function, perturbations of bioenergetic metabolism, specifically OXPHOS and glycolysis, that cannot be compensated for by the cell, will affect both the health and functioning of the cell. Here, we demonstrate how extracellular flux analysis that measures OCR, an indirect measurement of OXPHOS, and ECAR, an indirect measurement of glycolysis, gives a non-invasive, real-time, rapid insight into how anti-TB drugs affect the bioenergetic health of the cell, in the absence of cell death. Our data reveals the increased sensitivity of the extracellular flux analysis by the greater degree of variation in the response of the bioenergetic parameters of the anti-TB drug-treated cells in comparison to the % viability as the readout of the MTT assay (Fig. 6F and G). This strongly suggests that the bioenergetic parameters detect the effect of drugs on energy metabolism at much earlier timepoints prior to cell death, which is measured by the MTT viability assay. This was supported by correlation analysis in which the MTT assay had low, in some cases, negative correlations with the bioenergetic parameters (Fig. 6A-D), together with our PCA that demonstrated the MTT % Viability generated clustered separately from all the bioenergetic parameters (Fig. 6E).

Mitochondrial toxicity is now widely accepted as a common mechanism underlying drug induced organ toxicities (29, 31, 76), and is being increasingly detected in early stages of drug development using extracellular flux analysis (37, 38, 42). Mitochondrial toxicity of compounds is often assessed by growing the cells in the presence of high galactose (30, 42, 57), which forces the cells to use OXPHOS to produce ATP, thereby increasing the cells’ sensitivity to the effects of mitochondrial toxicants compared to cells grown in glucose (77, 78). However, the glucose-galactose switch does not sensitize all cell-types to mitochondrial cytotoxicity (79). Furthermore, in our study, we used glucose in our media to allow the cells to shift to glycolysis for ATP production (as evidenced by an increase in ECAR) should the drug adversely affect mitochondrial respiration (revealed by decreased OCR). Cells which cannot increase glycolysis in response to drug-impaired mitochondrial respiration, will not be able to meet the ATP requirements of the cell, thus increasing the cell’s susceptibility to adverse effects of the drug. Galactose does not allow this switch from OXPHOS to glycolysis. For this reason, we investigated the effects of the anti-TB drugs on hepatocytes, which rely more on OXPHOS, in addition to macrophages, which are more glycolytic, in the presence of glucose. Furthermore, we used the Cell Mito Stress test in the extracellular flux analysis as it has been demonstrated that the addition of the mitochondrial stressors, oligomycin, FCCP, antimycin A and rotenone, increased the sensitivity of the assay to detect mitochondrial dysfunction (36).

Addition of one of these stresses, FCCP, uncouples respiration from the production of ATP, resulting in maximal respiration (Max Resp) as a measure of the maximal activity of the electron transport chain (80). This enables the measurement of the SRC, which has been reported to be marker of cellular stress and mitochondrial dysfunction (81, 82). The SRC clustered separately from the other bioenergetic parameters in the hierarchical clustering of the effects of the anti-TB drugs on the bioenergetic parameters of the THP-1 cells (Fig. 3C), hMDMs (Fig. 3D) and when all three cell types were combined (Fig. 3A), demonstrating that treatment with the anti-TB drugs perturbs SRC differently to the other parameters. This is further supported by the correlation analyses of the different bioenergetic parameters, where the average Pearson’s correlation co-efficient was the lowest for SRC (Fig. 4A) in the THP-1 cells and hMDMs, and when the parameters of all three cell types were combined. Additionally, SRC also clustered separately from all the other bioenergetic parameters in our PCA of the bioenergetic parameters and the MTT % Viability (Fig. 6E), again reinforcing the distinctness of the SRC from the other bioenergetic parameters. Altogether, this demonstrates the high sensitivity of SRC to effects of the anti-TB drugs on the mitochondria. This has been supported by a study investigating the high throughput respirometry potential of the XF to detect mitochondrial biogenesis and toxicity (38). Known mitochondrial toxicants caused concentration-dependent depression in FCCP-uncoupled OCR with no significant decrease in the basal respiration of rabbit renal proximal tubule cells (RPTC). The authors concluded that the FCCP-uncoupled OCR can be used to uncover disrupted electron transport activity, and consequently mitochondrial damage, by toxicants even though basal metabolism is not impaired (38). As SRC is calculated from the response of OCR after addition of FCCP, our findings suggest that SRC can also be used to identify mitochondrial damage. Other studies investigating drug-induced cytotoxicity have also used FCCP-uncoupled OCR together with changes in initial OCR and ECAR in response to acute treatment or longer incubations with the drug on RPTC (78), and on RPTC and HepG2 (79) to specifically detect mitochondrial toxicity.

The choice of cell type to investigate cytotoxicity in drug development depends heavily on the environment, whether it be in industry or academia. Overall, we found the bioenergetic parameters of the macrophage models (THP-1 cells and hMDMs) were much more sensitive to the anti-TB drugs than the HepG2 cells (Table S1). This may be due to HepG2 cells having a greater SRC than the macrophages enabling them to tolerate further mitochondrial toxic insults than the macrophages with a lower SRC. This increased sensitivity is underscored by the distinct clustering of the effects of the of the anti-TB drugs on the bioenergetic parameters of the THP-1 cells, followed by broader groupings in hMDM cells, and the least distinction in the HepG2 cells (Fig. 5). These findings suggest that cytotoxic potential of new drug leads should not be investigated on one cell type alone. Although HepG2 cells give an indication of potential hepatotoxicity of early drug leads, macrophages are important in the innate immune response to *Mtb* infection and activation of the adaptive immune response. As the metabolism of the immune cells determines the immune functions of the immune cells, any alterations of the bioenergetic metabolism by new TB drug leads indicate the potential of these drugs to attenuate the immune response to *Mtb*. Our findings strongly advocate the use of two cell types for the screening of cytotoxicity in TB drug development: the HepG2 cells for detection of drug-induced liver toxicity and a macrophage model to detect the effects of the anti-TB drugs on immune cells, which are essential for control of infection.

Strikingly in all three cell types, the effects of EMB or MXF on the bioenergetic parameters of the cell types either group away or cluster together from the effects of the other anti-TB drugs on the bioenergetic parameters. These two drugs are not chemically related (Fig. S3) and do not have similar mechanisms of action on *Mtb*, which importantly, suggests that cytotoxic effects cannot be predicted from structure-activity relationships. EMB is thought to inhibit the biosynthesis of the cell wall, by inhibiting arabionsyltransferase required for the synthesis of arabinogalactan and lipoarabinomannan(83), whereas MXF is a fluoroquinolone that inhibits DNA gyrase that allows the untwisting required to synthesize two DNA helices from one DNA double helix (84). This is further supported by the effects of INH and RIF on the bioenergetic parameters clustering together in both the THP-1 and hMDM cells, although they are not chemically related nor have similar mechanisms of action. The clustering of the effects of EMB and MXF in addition to INH and RIF on the bioenergetic parameters of the hMDM cells were also mimicked in the hierarchical clustering of the effects of the same drugs on the levels of cytokines produced by hMDMs (Fig. 7). These findings of chemically unrelated drugs inducing similar effects on the bioenergetic parameters as well as similar patterns on hMDM cytokine production caution against associating cytotoxicity of new chemical entities with their chemical structures or mode of action against *Mtb*.

In conclusion, we have adopted real time extracellular flux analysis to detect early cytotoxic effects of the anti-TB drugs on the health of the cell prior to cell death by assessing the effects of the anti-TB drugs on the bioenergetic parameters of human HepG2 cells, THP-1 macrophages and hMDMs. In particular, SRC is the most sensitive measure of early mitochondrial toxicity induced by drug treatment. Interestingly, we found that chemically unrelated drugs with differing modes of action on *Mtb* cluster together in their similar effects on the bioenergetic metabolism of the cells, in particular the THP-1 cells. This points to the prudence of associating chemical structure and mode of action on *Mtb* with potential cytotoxicity patterns. Furthermore, our findings strongly advocate measuring the effects of new drug leads on the bioenergetic metabolism of macrophages in addition to HepG2 cells to assess cytotoxicity as this will not only assess hepatotoxicity but will also give an early indication of the potential of the new drug leads to modulate the immune functions of immune cells that might pose a risk to controlling *Mtb* infection. Thus, these findings can be used to establish a benchmark for cytotoxicity testing in future TB drug discovery.

## Methods

### Tissue Culture and differentiation

#### Human monocyte derived macrophages

Peripheral blood mononuclear cells (PBMCs) were isolated from buffy coats (South African National Blood Service). Briefly, 8 ml buffy coat was diluted in 27 ml Dulbecco’s phosphate buffered saline (DPBS) and overlaid onto 15 ml Histopaque® 1077. The buffy coat was separated (400 × *g*, 35 min, swing out bucket rotor; no acceleration, no brake). The PBMC-enriched layer was collected and washed with DPBS (1:1). PBMCs were pelleted (400 × *g*, 10 min) and washed in 50 ml DPBS (room temperature), then repeated with 50 ml DPBS (4°C). PBMCs were pelleted and resuspended in 5 ml separation buffer (DPBS, 2 mM EDTA, 0.5% (w/v) BSA, 4°C). CD14^+^ monocytes were isolated by magnetic cell sorting using MACS CD14-microbeads (Miltenyi, 130-505-201) according to manufacturer’s instructions. The monocytes were pelleted and resuspended in freezing solution (RPMI1640 with final concentrations of 10% (v/v) human serum, 1 mM sodium pyruvate, 10 mM HEPES, 1× non-essential amino acids, 2 mM GlutaMax™, 10% (v/v) DMSO). Monocytes were thawed in cell culture media (RPMI1640, 10% (v/v) human serum, 1 mM sodium pyruvate, 10 mM HEPES, 1× non-essential amino acids, 2 mM GlutaMax™) counted using trypan blue to assess the viability and seeded directly into XFe96 cell microtiter plates at a density of 8×10^4^ cells per well in a volume of 80 μl. The monocytes were terminally differentiated into macrophages with 100 ng/ml GM-CSF for 6 days, with a media change (including the GM-CSF) on day 4. On the sixth day, the macrophages were treated with the anti-TB drugs for 24 hrs prior to extracellular flux analysis on the XFe96 and viability analysis using the MTT assay.

#### THP-1 macrophages

THP-1 monocytes (ATCC TIB-202) were cultured in RPMI1640 (final concentrations: 10% (v/v) FBS, 25 mM D-glucose, 10 mM HEPES, 1 mM sodium pyruvate, 2 mM L-Glutamax, 0.05 mM β-mercaptoethanol) under standard tissue culture conditions (37°C, 5% CO_2_). Cells were washed in fresh media, counted, and seeded in the XFe96 cell culture plate at a density of 100 000 cells per well in 80 μl RPMI1640 culture media and terminally differentiated with 25 nM phorbol 12-myristate-13-acetate (PMA) for 3 days. On the fourth day, fresh media without PMA was supplied to the cells, and on the fifth day the cells were treated with the anti-TB drugs for 24 hrs.

#### HepG2 cells

HepG2 cells (ATCC HB-8065) were cultured in DMEM supplemented with 10% (v/v) FBS. To seed, cells were washed with warm DPBS and lifted with warm 1× Trypsin-DPBS. Trypsin was deactivated with the addition of culture media. Cells were harvested, pelleted (400 × *g*, 5 min), resuspended in fresh media and seeded at a density of 25 000 cells per well in 80 μl DMEM culture media of the XFe96 cell microtiter plate. Cells adhered naturally overnight, followed by 24 hrs treatment with the anti-TB drugs.

### Anti-TB drug treatment and Agilent Seahorse Cell Mito Stress Test (CMST)

Stock solutions of anti-TB drugs were prepared in DMSO or DPBS where possible (Table 1). Working drug solutions were prepared in the respective media, and the final concentration of DMSO per well did not exceed 0.2% (v/v), except for the highest concentrations (50x and 100x MIC) of clofazimine and linezolid (0.5% and 1% (v/v)), in which case 0.5% and 1% DMSO controls were included in the assays. Following seeding and/or differentiation of each cell type, the supernatant was aspirated, and the cells were treated with four concentrations of each anti-TB drug: 1x, 10x, 50x and 100x the MIC values in Table 1 in 8 replicates for 24 hrs in a total volume of 80 μl/well. Cells were also treated with two drug combinations: (1) INH, RIF, PZA, EMB and STR, or (2) INH & RIF at 1x and 10x MIC of all the drugs in the combination for 24 hrs. The following day, the cells were washed twice with CMST media (DMEM, 30 mM NaCl, 5 mM HEPES, 2 mM L-Glutamax, 1 mM sodium pyruvate, pH 7.4) and the final volume was brought up to 180 μl with CMST media. The media in the cell plate was degassed for a minimum of 30 min in a non-CO2 incubator. The mitochondrial modulators oligomycin, carbonyl cyanide-4 (trifluoromethoxy) phenylhydrazone (FCCP), rotenone and antimycin A were prepared in CMST media from DMSO stocks at 10x the concentrations given in Table 2. The pH of the solutions was adjusted to 7.4 at 37°C and loaded into the ports of the XFe96 cartridge as indicated in Table 2 (85, 86). The extracellular flux of the cells was analyzed on an XFe96 using the Cell Mito Stress Test (CMST) protocol with 3 minutes of mixing and 4-minute measurements.

**Table 1.**
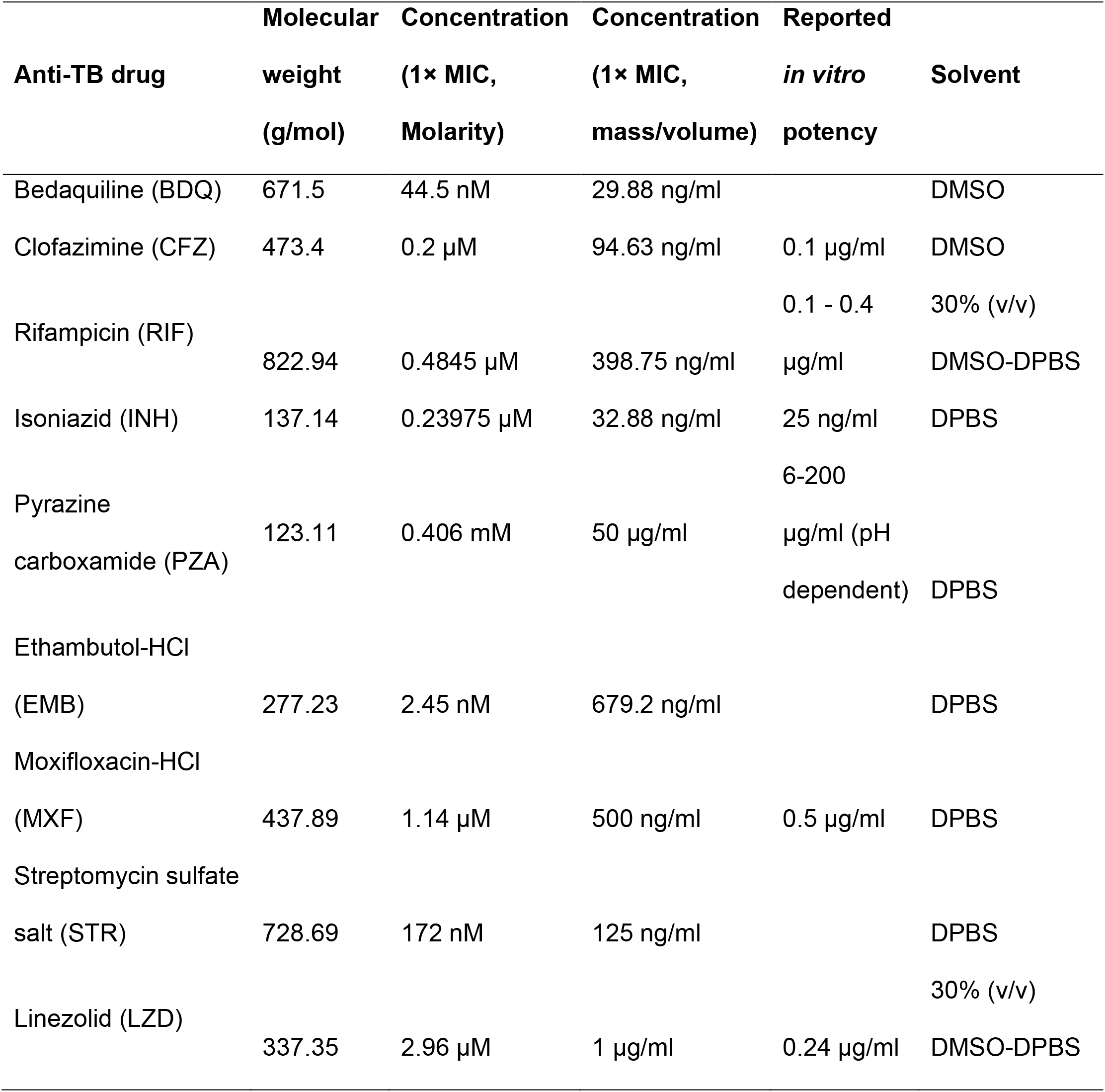
Concentrations and preparation of Anti-TB drugs.

**Table 2:** Final concentrations of mitochondrial modulators used for CMST.

| Modulator | Port* | Volume (μl)<sup>#</sup> | Concentration (μM) |  |  |
| --- | --- | --- | --- | --- | --- |
|  |  |  | hMDM | THP-1 | HepG2 |
| Oligomycin | A | 20 | 1.5 | 1.5 | 3 |
| FCCP | B | 22.5 | 1 | 1 | 2.5 |
| Rotenone and Antimycin A | C | 25 | 2.5 | 0.5 | 0.5 |

\* Port on XFe96 cartridge.
<sup>#</sup>Volume loaded into the port

### Normalization of extracellular flux by protein concentration

Following XFe96 analysis, the supernatants were aspirated from all the wells leaving behind approximately 10 μl of the supernatant in each well, and the cells were fixed with the addition of 10 μl formalin/well. The cells in each well were lysed by adding 20 μl 25 mM NaOH/well. BSA standards (5 μl) were added to the control wells without cells (Lanes 1 and 12) ranging from 0.125 – 2 mg/ml (BioRad 500-0202) and treated with formalin and NaOH at the same concentrations as the cells. Bradford reagent (150 μl, BioRad 500-0205) were added to all the wells, and the plate was incubated in the dark for 5 min. The absorbance of each well was measured at 595 nm using a Biotek Synergy H4 Hybrid spectrophotometer and the standard curve generated from the BSA standards in lanes 1 and 12 was used to calculate the protein concentrations of each well. These protein concentrations were used to normalize the bioenergetic parameter data (OCR and ECAR) were normalized using the protein concentration in the Agilent Seahorse Wave desktop software (version 2.6). The CMST assay parameters were calculated using the Agilent Seahorse Biosciences Cell Mito Stress Test Report Generator.

### MTT viability assay

To assess the viability of the cells under the identical conditions used for the extracellular flux analysis, the cells were seeded into XFe96 cell microtiter plates and treated as for the XFe96 assay. Media in lanes 1 and 12 of the microtiter plate served as the negative controls. The MTT reagent (3-(4,5-dimethylthiazol-2-yl)-2,5-diphenyltetrazolium bromide; Invitrogen M6494) was prepared and the assay was performed according to manufacturer’s instructions. Briefly, after overnight drug treatments, the supernatant was aspirated, leaving 25 μl of supernatant behind to avoid lifting the cells. The volume was bought up to 100 μl with the appropriate media, and 10 μl of MTT reagent (5 mg/ml in DPBS) was added to each well, and incubated for 4 hours (37°C, 5% CO_2_). Supernatant was aspirated and the formazan crystals was dissolved with 50 μl DMSO in each well followed by 10 min incubation at room temperature. The absorbance of each well at 540 nm was measured using the Biotek Synergy after mixing by trituration. The percentage viability of the cells was calculated as follows:

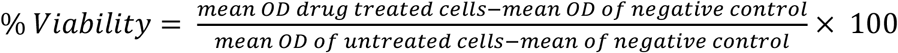

### Cytokine measurements in the culture supernatant fluid

Culture supernatant was collected from the hMDM cells treated with the anti-TB drugs for 24 hrs prior to XF runs and stored at −80 °C. The cytokine levels were measured using the magnetic bead-based Bio-Plex Pro Human cytokine 27-Plex (Bio-Rad) according to manufacturer’s instructions and measured the cytokines using the Bio-Plex 200 instrument. Using the Bio-Plex Manager Software and standard curves of each cytokine, the concentrations of the cytokines (pg/ml) were calculated from the median fluorescence intensity (MFI). Four replicates were used for each concentration of each anti-TB drug analyzed.

### Statistical analyses

Two-way ANOVA was performed using GraphPad Prism for the bioenergetic parameters. Principle components analyses (PCA), Hierarchical clustering (heatmap), and Pearson’s correlation were performed using software package Partek Genomic Suite (PGS, Partek, MO, US. Partek.com) according to factory settings and user manual. P-values less than 0.05 was considered significant. Briefly, z-normalization was performed before hierarchical clustering. Euclidean dissimilar matrix and average linkage similarity were used. In PCA analysis, all variables have assumed having equal influence on principle components (PC).

## Acknowledgements

This work was supported by NIH Grants R01AI134810, R01AI137043, R01AI152110, R33AI138280, a Bill and Melinda Gates Foundation Award (OPP1130017), the South African (SA) Medical Research Council and a SA NRF BRICS Multilateral grant to A.J.C.S.

